# Differential impact on motility and biofilm dispersal of closely related phosphodiesterases in *Pseudomonas aeruginosa*

**DOI:** 10.1101/697938

**Authors:** Yu-Ming Cai, Andrew Hutchin, Jack Craddock, Martin A. Walsh, Jeremy Stephen Webb, Ivo Tews

## Abstract

Bacteria typically occur either as free-swimming planktonic cells or within a sessile, biofilm mode of growth. In *Pseudomonas aeruginosa*, the transition between these lifestyles is known to be modulated by the intracellular secondary messenger cyclic dimeric-GMP (c-di-GMP). We are interested in the control of distinct biofilm-relevant phenotypes in *P. aeruginosa* through the modulation of intracellular c-di-GMP. Here, we characterise motility and associated biofilm formation and dispersal in two pairs of related multi-domain proteins with putative c-di-GMP turnover domains, selected to contain additional PAS (Per-Arnt-Sim) homology domains known for their ability to process environmental stimuli. The enzymes PA0861 (RbdA) and PA2072 have distinct functions despite their similar domain structures. The Δ*rbdA* deletion mutant showed significantly increased biofilm formation while biofilm formation was impaired in Δ*PA2072*. Using a GFP transcriptional reporter fused to the cyclic di-GMP-responsive *cdrA* promoter, we show correlation between biofilm phenotype and c-di-GMP levels. Both proteins are shown to play a role in nitric oxide (NO) induced biofilm dispersal. We further studied pseudo-enzymes of similar architecture. PA5017 (DipA) is an inactive cyclase, and PA4959 (FimX) is described here as an inactive cyclase/phosphodiesterase. Loss of swimming and twitching motilities, respectively, is observed in deletion variants, which correlated with NO-induced biofilm dispersal phenotypes, as Δ*dipA* dispersed less well while Δ*fimX* dispersed better than wild type. The study highlights how *Pseudomonas* differentiates c-di-GMP output – in this case motility – using structurally very similar proteins and underlines a significant role for pseudo-enzymes in motility regulation and associated biofilm dispersal.

**Importance:** Bacterial biofilms exert pervasive economic and societal impact across a range of environmental, engineered and clinical contexts. The secondary messenger cyclic guanosine di-phosphate, c-di-GMP, is known to control the ability of many bacteria to form biofilms. The opportunistic human pathogen *Pseudomonas aeruginosa* PAO1 has 38 putative enzymes that can regulate c-di-GMP turnover, and these proteins modulate various cellular functions and influence bacterial lifestyle. The specific protein sensory domains and mechanisms of motility that lead to biofilm dispersal remain to be fully understood. Here we studied multi-domain proteins with the PAS (Per-Arnt-Sim) homology domains, these being classic sensors to environmental stimuli. Our study demonstrates the significant roles for the pseudo-enzymes PA4959 (FimX) and PA5017 (DipA) in regulation of biofilm phenotype and motility. Further, enzymes with highly homologous structures, such as PA0861 (RbdA) and PA2072, have almost orthogonal function in biofilm and motility control.

## Introduction

*Pseudomonas aeruginosa* is a gram-negative bacterium known for its environmental versatility. As an opportunistc pathogen, *P. aeruginosa* causes disease, particularly in immune compromised individuals, and is a major cause of morbidity and mortality in cystic fibrosis (CF) patients with chronic colonisation in lungs and airways^1^. The ability of *P. aeruginosa* to form biofilms within CF patients causes increased antibiotic tolerance, which makes treatment of infections problematic in clinical settings^2^.

*P. aeruginosa* biofilm formation and dispersal are known to correlate with intracellular concentrations of the secondary messenger, cyclic dimeric-GMP^3,4^. The production and degradation of c-di-GMP relies on two enzymatic activities. Diguanylate cyclases (DGCs) synthesise c-di-GMP from two GTP molecules, while phosphodiesterases (PDEs) hydrolyse the secondary messenger to linear pGpG, and in some cases GMP^3^. PAO1 encodes 17 different proteins with a DGC domain, 5 with a PDE domain, and 16 that contain both of these domains as a fusion (DGC-PDE)^5^. Within *P. aeruginosa*, intracellular c-di-GMP levels can be regulated by a number of environmental cues, including nutrient availability and the presence of nitric oxide^6,7^. Indeed, DGC and PDE-containing proteins typically also have one or several putative regulatory domains, such as the ubiquitous PAS (Per-Arnt-Sim) domain^8^.

In *P. aeruginosa* PAO1, twelve genes code for proteins with PAS domains linked to DGC domains, nine of which additionally contain PDE domains^5^, namely *pa0285, pa0290, pa0338, pa0575*^9^, *pa0847, pa0861* (*rbdA*^10^), *pa1181* (*yegE*), *pa2072*, *pa4601* (*morA*^11–13^), *pa4959* (*fimX*^14^), *pa5017* (*dipA*^15^), and *pa5442*. The recurrence of proteins with similar architecture, and which play a role in biofilm regulation, poses two key questions: Firstly, if DGCs and PDEs regulate the transition between sessility and the planktonic state, how do the specific protein sensory domains then control mechanisms of motility? Secondly, while speculated to be c-di-GMP sensors, what is the role of proteins with DGC or PDE domains that are rendered inactive due to mutations altering the active sites of enzymes, and thus their catalytic activity? This knowledge is urgently needed to understand the mechanisms regulating biofilm dispersal.

The main motility types in *P. aeruginosa* PAO1 are flagella mediated swimming; pili mediated twitching; and the complex swarming motility, which relies on flagella, pili and surfactants and involves multicellular group movement on a surface^16,17^. The regulatory relationships between c-di-GMP and different motility types have been investigated for a number of *P. aeruginosa* proteins. CheA and the partitioning protein ParP were shown to control the subcellular localisation of DipA in chemotaxis, controlling reduction of c-di-GMP and swimming motility in PAO1^18^. Deletion of RbdA was also reported to reduce swimming^10^. Over-expression of diguanylate cyclases can inhibit swimming motility significantly, as seen with the DGC SadC^19^. SadC is part of a regulatory circuit involving the phosphodiesterase BifA that regulates swarming through modulation of flagellar reversal rates^20,21^, while the flagellar stators MotAB and MotCD directly respond to c-di-GMP levels^22^. Most studies of swarming have focussed on the role of this motility in biofilm formation^20,23,24^, but a time-lapse microscopy study reported a role for swarming in dispersal^25,26^. A bidirectional relationship between c-di-GMP and swarming through so called Mot proteins is established for *P. aeruginosa* PA14^27^, however data for *P. aeruginosa* PAO1 are sparse.

To dissect their roles in biofilm regulation, we created *P. aeruginosa* PAO1 deletion mutants of the twelve identified PAS-DGC-PDE proteins and compared them to wild-type PAO1. We selected the highly homologous enzymes PA0861 (RbdA) and PA2072 which we show to have different roles in biofilm control, the protein PA5017 (DipA) that is a phosphodiesterase coupled to an inactive diguanylate cyclase^15^, and the protein PA4959 (FimX) with an inactive cyclase domain and a phosphodiesterase domain with contradicting reports of catalytic activity^14,28–31^ that are addressed here. Deletion mutants were characaterised for biofilm morphology, EPS production, c-di-GMP level as quantified using a GFP reporter system, and variation in motility. Dispersal phenotype for each deletion mutant was also assessed using the nitric oxide (NO) donor S150. Our data reveal specific alterations in phenotype and motility that play significant roles in both biofilm structure and in the response to NO which correlate with NO-induced swarming motility and biofilm dispersal.

## Materials and methods

### Bioinformatics and homology modelling

To assign domain architectures, the EMBL SMART web server was used (http://smart.embl-heidelberg.de^32^). To predict catalytic activities, amino acid sequences of individual catalytic domains were aligned against equivalent domains known to be catalytically active and for which a structure has been deposited in the PDB, using the clustal web server (https://www.ebi.ac.uk/Tools/msa/clustalo/)^33^. The data were complemented through protein structure homology modelling carried out using the SWISS-MODEL server (https://swissmodel.expasy.org)^34^. As a template for DGC domains, PleD (PDB 2V0N^35^) was used, while MorA (PDB 4RNH^11^) was used as a template for PDE domains. While inspecting substrate binding sites and catalytic centres of the homology models, conservation of catalytic residues and compatibility with substrate binding was taken as an indicator of enzymatic activity.

### Bacterial strains and culture media

Bacterial strains and plasmids used in this study are listed in **Table S1**. Routine overnight cultures were grown in lysogeny broth (LB) medium. Biofilms were grown in standard M9 minimal medium. Antibiotics were used at the following concentrations: for *P. aeruginosa* PAO1, gentamicin was used at 30 μg/ml, carbenicillin at 400 μg/ml, kanamycin at 300 μg/ml and tetracycline at 60 μg/ml; for *E. coli* S17-1, gentamicin was used at 15 μg/ml, ampicillin at 100 μg/ml, tetracycline at 30 μg/ml, kanamycin at 50 μg/ml and streptomycin at 50 μg/ml.

### Isogenic P. aeruginosa PAO1 mutants

Isogenic mutants were constructed by replacing the coding regions of each gene with a gentamicin resistance cassette as previously described^36^. The Gm cassette was amplified from pPS856^37^ using primers Gm-F and Gm-R (primer sequences listed in **Table S2**). For each gene, flanking up- and downstream regions (approximately 350-400 bp) were amplified by standard PCR, digested with EcoRI and HindIII, and ligated to the amplified Gm cassette. To generate KO plasmids, the PA-up-Gm-PA-dn fragments were inserted into the SmaI site of the pEX100T suicide vector, which contains the sacB gene for counter selection and an ampicillin resistance gene on the backbone. Plasmids were introduced into *E. coli* S17-1 by chemical transformation and then transferred into PAO1 by conjugation. Transconjugants were first selected on *Pseudomonas* isolation agar containing 30 μg/ml Gm and then patched onto LB agar with 10 % sucrose and 30 μg/ml Gm and LB agar with 400 μg/ml carbenicillin. Colonies that only grew on LB agar with 10 % sucrose and 30 μg/ml Gm but not on LB agar with 400 μg/ml carbenicillin were selected as double-recombinant mutants and confirmed by PCR and sequencing. For the generation of the double Δ*pilAfliM* mutant, the gentamycin cassette was used for the introduction of a single mutant, with a kanamycin cassette used to introduce the second mutation. This Km cassette was amplified from pCR™4-TOPO™ (Invitrogen) using primers Km-F and Km-R.

### Batch culture P. aeruginosa PAO1 biofilms

Overnight cultures were diluted into fresh M9 media (OD_600nm_ ~0.01) to inoculate microtiter plates or MatTek plates (P35G-1.5-14-C), using 100 μl or 3 ml of diluted culture, respectively. Microtiter plates were incubated statically, while MatTek plates were shaken at 50 rpm to create shear force facilitating biofilm formation, changing M9 media every 24 hrs. Biofilms in microtiter plates were stained with 0.1 % (w/v) crystal violet and dissolved in 30 % (v/v) acetic acid. Biofilms in MatTek plates were stained with LIVE/DEAD^®^ BacLight (Invitrogen) and examined by confocal laser scanning microscopy. Crystal violet staining was determined at a wavelengths of 584 nm, while wavelength of 488 nm and 561 nm were used for SYTO-9 and propidium iodide excitation, respectively. At least 3 image stacks were taken from random locations in each MatTek plate. Biofilm biomass and surface coverage were analysed by COMSTAT^38^, while microcolony sizes were analysed by merging confocal images stacks with ImageJ to measure identified microcolonies with DAIME^39^. For NO donor experiments, 250 μM Spermine NONOate (S150, Sigma-Aldrich) was added into MatTek biofilm plates and incubated at 37 °C for 2 hrs to trigger dispersal.

### NO donor release determination

The concentration of NO donor released from specific NO donors (**Table S3**) was quantified using a chemiluminescence-based method adapted from Piknova *et al*.^40^. Once released from the donor, free NO and O_3_ formed activated NO_2_, which emits a detectable photon upon relaxation. Both SNP and S150 were dissolved in M9 medium and tested at 37 °C. For SNP, two concentrations were tested under different light conditions – 5 μM was tested under normal light while 500 mM was tested with/without a cold light trigger. For S150, 5 μM was tested under normal light. Values were compared with standards determined from a sodium nitrate, tri-iodine solution. Details of the method used are found in the supplementary data.

### Determination of swarming motility

Swarming agar plates were prepared from 0.5 % (w/v) agar (Sigma Aldrich) in 8 g/L nutrient broth (Oxoid) and 5 g/L glucose (Sigma Aldrich). NO donor swarming plates additionally contained SNP at a final concentration of 1 μM. Swarming plates were dried under laminar flow for 40 mins and inoculated with 3 μl late exponential culture. Plates were incubated at 37 °C for 24 hrs under normal laboratory light conditions. For swarming agar with SNP, 10 mM SNP stock solution was made in sterilised PBS and then incorporated into 50 °C swarming agar to a final concentration of 1 μM.

### Determination of swimming motility

Tryptone broth (10 g/L tryptone, Oxoid) with 5 g/L NaCl and 0.3 % (w/v) agarose (Melford) were used to prepare agar plates for swimming motility assays. The plates were dried under laminar flow for 15 mins and inoculated from an overnight LB agar plate with a sterile 2 μl pipette tip. Incubation at 30 °C was carried out for 20 hrs.

### Determination of twitching motility

Twitching agar plates were prepared from LB broth (Miller) with 1 % (w/v) agar (Sigma Aldrich). Plates were dried under laminar flow for 1 hr and inoculated form an overnight LB agar plate by stabbing with a sterile toothpick through the agar to the bottom of the plate. Plates were incubated at 37 °C for 24 hrs.

### EPS extraction and quantification

Biofilms with an initial inoculum of diluted overnight culture (OD_600nm_ ~0.01) were cultured in tissue-culture treated dishes (Corning, UK, D×H 100mm×20mm). Cell scrapers were used to harvest bacteria, re-suspending and vortexing them in 3 ml PBS, from which 100 μl were taken for CFU counts. Supernatants were directly analysed for soluble polysaccharide and protein content, while 2 ml 0.85 % NaCl and 12 μl 37 % formaldehyde (Sigma-Aldrich, UK) were added to the cell pellets after centrifugation to determine the insoluble polysaccharide and protein content. Mixtures were vortexed and incubated at 4 °C for 1 hr. Addition of 2 ml 1 M NaOH and 0.5 ml H_2_O was followed by further incubation for 3.5 hrs. Supernatants were harvested by centrifugation at 4000×*g* (40 mins, 4 °C) and freeze-dried. Dried supernatant was dissolved in water and adjusted to pH 7 using H_2_SO_4_. The polysaccharide content was determined using the phenol-H_2_SO_4_ method^41^ by measuring the absorbance at 492 nm; with glucose used as a standard. The protein content was determined using the Coomassie (Bradford) protein assay kit (Thermo Scientific), measuring absorbance at 595 nm; calibrated to a standard of bovine serum albumin.

### *Determination of the relative level of c-di-GMP in vivo* (adapted from Rybtke *et al*^42^)

The c-di-GMP reporter plasmid (courtesy of A. Filloux, Imperial College London, UK) was introduced into *P. aeruginosa* by conjugation. Cultures were inoculated in 10 ml M9 with 60 μg/ml tetracycline (OD_600nm_ 0.001) and incubated for 22 hrs at 37 °C by shaking at 180 rpm. Strains without reporter were used as a negative control. For NO donor treatment, 25 μM S150 was added to cultures before incubation for an additional 2 hrs. Bacterial cultures (100 μl) were transferred into black polystyrene flat bottom Greiner CELLSTAR^®^ 96 well plates to determine fluorescence intensity, and into clear polystyrene flat bottom CELLSTAR^®^ 96 well plates to determine cell density. Arbitrary fluorescence intensity units (FIU) were determined using a 485 nm sharp-cut excitation filter and a 520 nm sharp-cut emission filter with a gain of 1500 on a BMG LABTECH FLUOSTAR plate reader. Relative fluorescence units (RFU) were determined from the FIU normalised by cell density (OD)^43^:

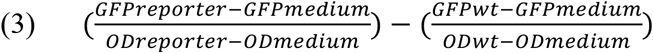

## Results

### Structure and enzymatic activity of twelve PAS-DGC-PDE proteins

The *Pseudomonas aeruginosa* PAO1 genome encodes 38 proteins with either diguanylate cyclase (DGC) or phosphodiesterase (PDE) domains. These are often found in multi-domain proteins with a variety of auxiliary domains, such as putative regulatory domains or membrane segments. Within PAO1, additional domains are exclusively found N-terminal to the DGC or the PDE domains, or to the DGC-PDE tandem^5^. While membrane domains are most common, the PAS domain, named after the Per-Arnt-Sim homology^8^, is the second most commonly found domain. To better understand the widespread usage of this architecture in DGC and PDE proteins, we studied the multi-domain proteins that contain PAS sensor domains together with DGC and/or PDE domains.

Twelve proteins were identified in *P. aeruginosa* PAO1 that contained PAS domains together with DGC and/or PDE domains, **Figure 1**. Of these, eight are associated to the membrane, as evidenced by the presence of at least one predicted transmembrane segment. The number of predicted PAS domains in these proteins varies from one to four. Further sensory or regulatory domains are predicted for several of these proteins. For example PA0575 also contains a periplasmic domain, PA5017 (DipA) carries an additional GAF domain, PA0847 contains a HAMP domain and PA4959 (FimX) contains a REC domain.

All twelve proteins contain DGC domains, with nine of them containing an additional PDE domain. For several of these proteins, enzymatic activity has been reported; others have been shown to be inactive as is indicated in solid colour in **Figure 1**. For the proteins that were not reported to be either active or inactive, we have investigated conservation of the enzymatic signature motifs as an indication of catalytic activity. For DGC domains, conservation of the GGDEF motif is required, as the aspartate and glutamate residues in this motif are required for catalysis. For EAL domains, the EAL and DDFGTG motifs are required, contributing a glutamate and two aspartates to the catalytic centre, respectively.

We have investigated the conservation of these motifs through sequence alignment^33^. Where present, deviations from archetypal catalytic motifs were compared to existing examples of identical atypical motifs and analysed using homology modelling. The potential for substrate binding was ascertained from the homology models by inspection of catalytic centres. DGC domains were modelled after the substrate-bound structure of *Caulobacter crescentus* PleD (PDB 2V0N^35^), and PDE domains were modelled after the substrate-bound structure of *P. aeruginosa* MorA (PDB 4RNH^11^). Domains with predicted enzymatic activity are shown in striped shading in **Figure 1**.

Of the twelve PAS-DGC and PAS-DGC-PDE domain proteins, two have been identified to contain inactive c-di-GMP metabolising domains. PA4959 (FimX) has been experimentally determined to have a degenerate DGC domain, and while enzymatic activity of the PDE domain remains a somewhat contested subject within the field due to contrasting reports of c-di-GMP hydrolysis, with those reporting activity doing so at a very low level, we have labelled the domain as inactive, in line with others, as it contains degenerate catalytic motifs not compatible with the established mechanism of EAL catalysed PDE activity^14,28–31^. PA5017 (DipA) contains an active PDE, but an inactive DGC^15^.

### The deletion mutants of Δ*fimX*, Δ*dipA*, Δ*rbdA* and Δ*pa2072* have characteristic biofilm phenotypes

Deletion mutants for the twelve PAS domain containing DGC/PDE proteins shown in **Figure 1** were constructed as described in the Methods. We selected the two mutants, Δ*dipA* and Δ*fimX*, to understand how (partially/fully) inactive sensor-enzymes function in biofilm and motility modulation. Additionally, we selected the mutants Δ*rbdA* and Δ*pa2072* to characterise through which trait they exert effects on biofilm behaviour, as they showed a striking difference in biofilm phenotype from an initial phenotypical screen (data not shown). These genes encode proteins that are highly homologous in structure.

PAO1 biofilms reached maturation stages with substantial microcolonies after 48 hrs and maximum surface coverage after 72 hrs under the experimental conditions used in this study. To highlight differences in biofilm formation, biofilms were inspected after 48 hrs. Upon a first visual inspection of CSLM images, **Figure 2A**, much thicker biofilms than PAO1 WT were observed in Δ*rbdA*, with increased microcolony formation but equal height at the base of the biofilm. The opposite was true for Δ*pa2072* that lacks substantial microcolony formation. The biofilms of the mutants Δ*dipA* and Δ*fimX*, on the other hand, are similar to PAO1 WT.

**Figure 1.** *Pseudomonas aeruginosa* proteins with PAS and DGC/PDE domains. The domain structure is shown on the right as determined by the SMART domain prediction server, with the sequence length as indicated. The observed sequence of the signature motifs for DGC domains (GGDEF) and PDE domains (EAL, DDFGTG) is given. Where enzymatic activity has been determined experimentally, this is indicated in green (or red, for no activity). Sequence alignment and homology modelling were used to predict enzymatic activity for the remaining proteins, as indicated by shading.

**Figure 2.**
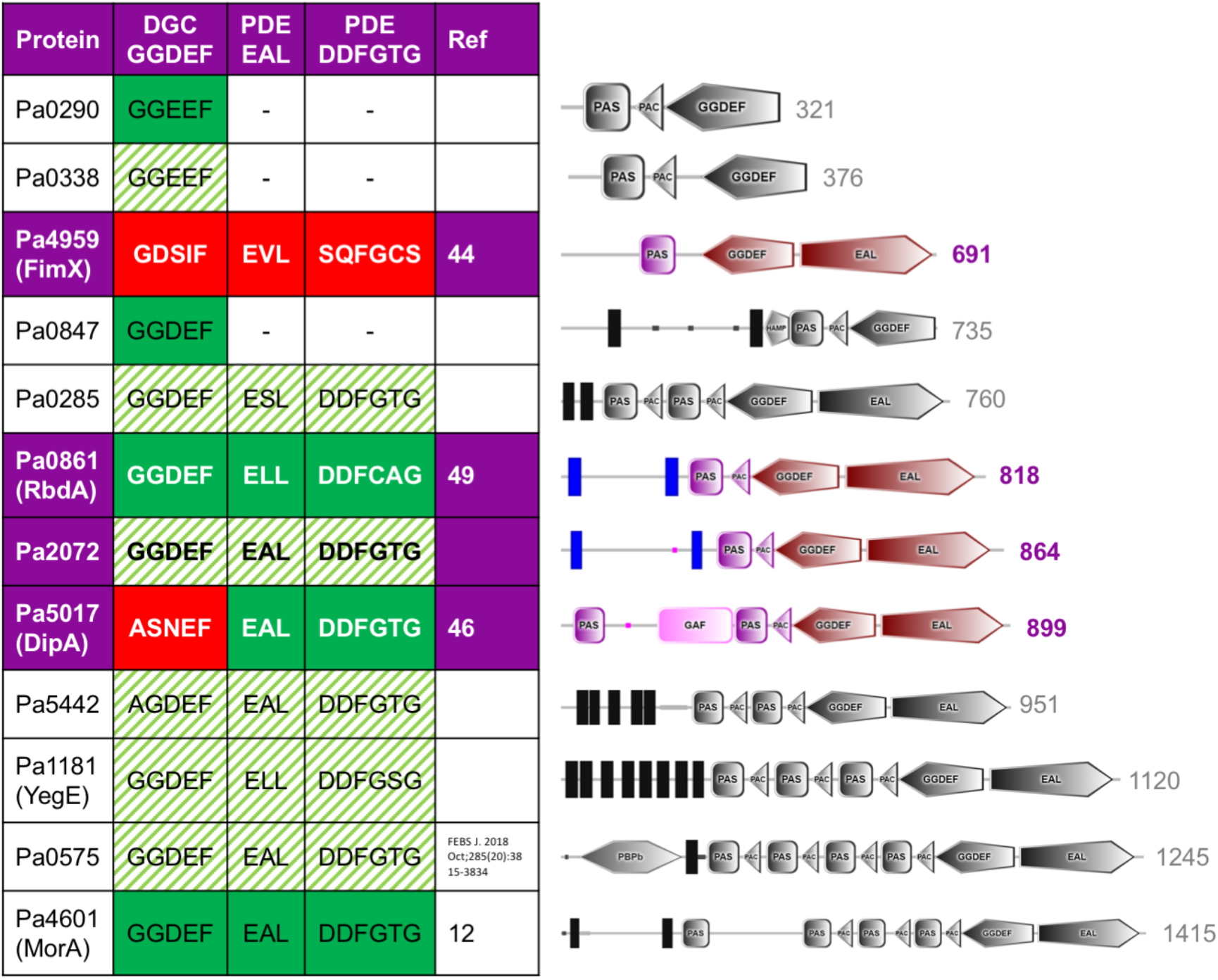
Biofilm phenotypes of deletion mutants. Biofilms formed in MatTek plates after 48 hrs for Δ*rbdA*, Δ*pa2072*, Δ*fimX* and Δ*dipA*, compared with PAO1 WT. (A) Confocal laser scanning micrographs at 63x magnification (scale bar 25 μm); live cells are stained with SYTO-9 (green); dead cells are stained with propidium iodide (red). (B) Quantification of microcolony size. (C) Quantification of biofilm biomass. (D) Quantification of surface coverage. For data shown in B-D, the Welch T-test was used to determine significances, where *** denotes a confidence level of p<0.01. Data acquired from 3 independent experiments.

The biomass, the surface coverage and the maximum microcolony sizes were determined to distinguish the development of morphologies. The data, normalised to PAO1 WT, are shown in **Figure 2B-D**. The different appearance in the thicker biofilms of Δ*rbdA* is reflected in a nearly five-fold increase of microcolony size, **Figure 2B**, and a more than two-fold increase in biomass, **Figure 2C**. In contrast, Δ*pa2072* showed much reduced microcolony size compared to WT, and only one fifth of the biomass. These phenotypes also affected biofilm surface coverage, which for Δ*pa2072* was only about one third of WT **Figure 2D**. The data suggest that PA2072 and RbdA play opposite roles in PAO1 biofilm formation, which is surprising given their very similar domain architecture, **Figure 1**.

Interestingly, this analysis reveals also a difference for the less conspicuous Δ*fimX* deletion mutant, as the biofilms show half of the WT biomass, **Figure 2C**, combined with unchanged maximum microcolony size or surface coverage, **Figure 2B** and **2D**, suggesting a different biofilm structure for Δ*fimX*. Similarly, Δ*dipA* shows comparable biomass and microcolony size compared with WT, **Figure 2B** and **2C**, but increased surface coverage, **Figure 2D**, again pointing at subtle differences in biofilm structure.

The protein and polysaccharide content of the EPS matrix, normalised to cell density, was determined. Consistent with the increased biomass and surface coverage observed for the deletion mutant Δ*rbdA*, **Figure 2C** and **Figure 2D**, enhanced total protein and polysaccharide are observed, **Figure 3**. The structurally related protein Δ*pa2072*, however, produced comparable EPS with WT, again pointing at functional differences between these two homologous proteins. The increased protein and polysaccharide mass of Δ*fimX* does not directly correlate to biofilm structure but is another indicator of a difference in the structure of Δ*fimX* biofilms.

**Figure 3.**
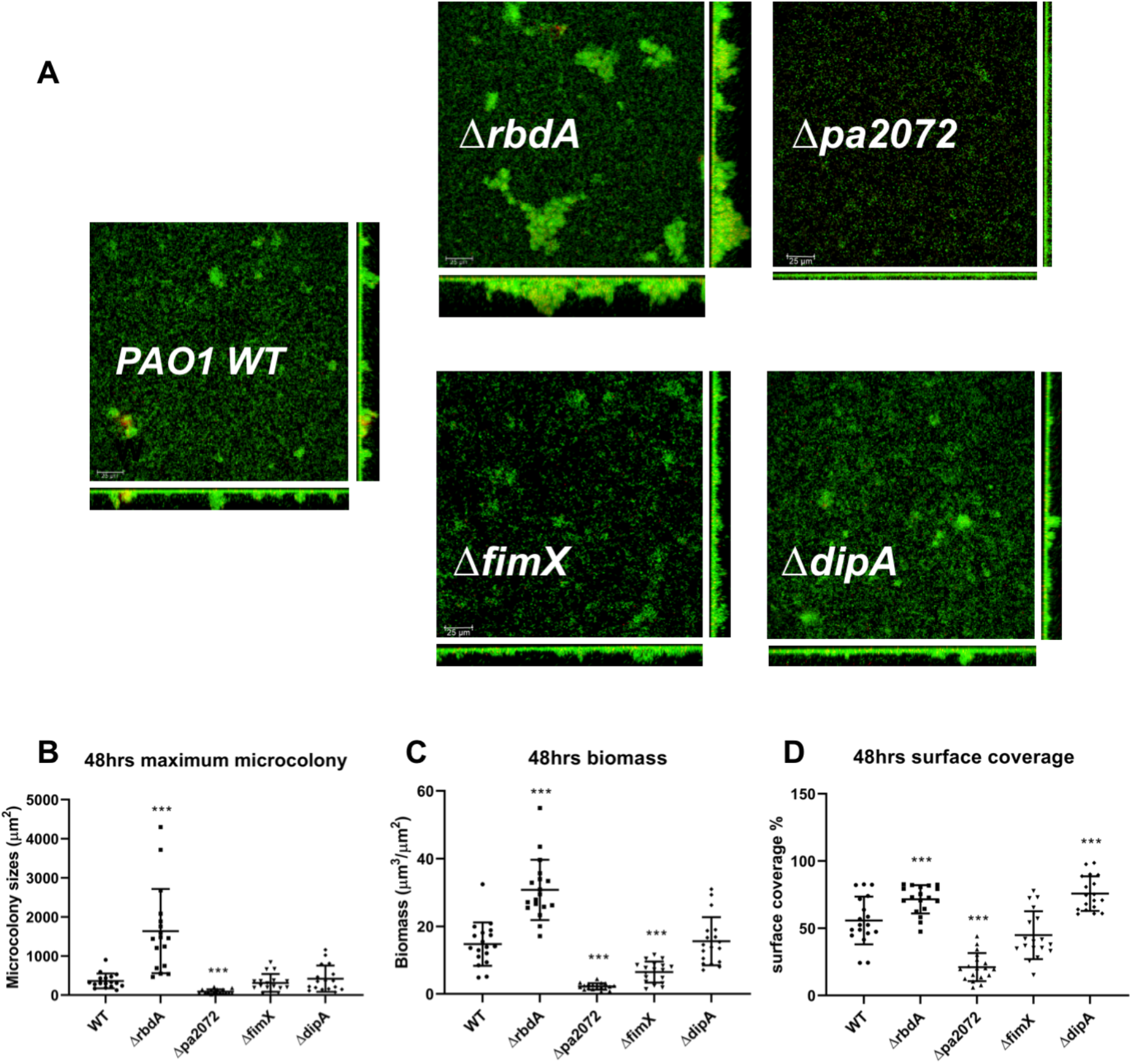
EPS total protein and polysaccharide. Quantification of EPS from 10^8^ cells of 48 hrs biofilms of Δ*rbdA*, Δ*fimX*, Δ*dipA*, and Δ*pa2072*, compared with PAO1 WT. Total protein and polysaccharide mass are shown as comparison to PAO1 WT, where the Welch T-test was used to determine the significances, and *** denotes a confidence level of p<0.01. Data acquired from 3 independent experiments.

The differences of the Δ*dipA* and *ΔfimX* deletion variants to PAO1 WT are aparent when bacterial motility is investigated. Flagellum mediated swimming motility is severely supressed in Δ*dipA*, and the phenotype is indeed similar to the flagellum mutant Δ*fliM* and the pili flagellum double mutant Δ*pilA*Δ*fliM*, **Figure 4A**. This phenotype is similar to the *ΔrbdA* mutant.

**Figure 4.**
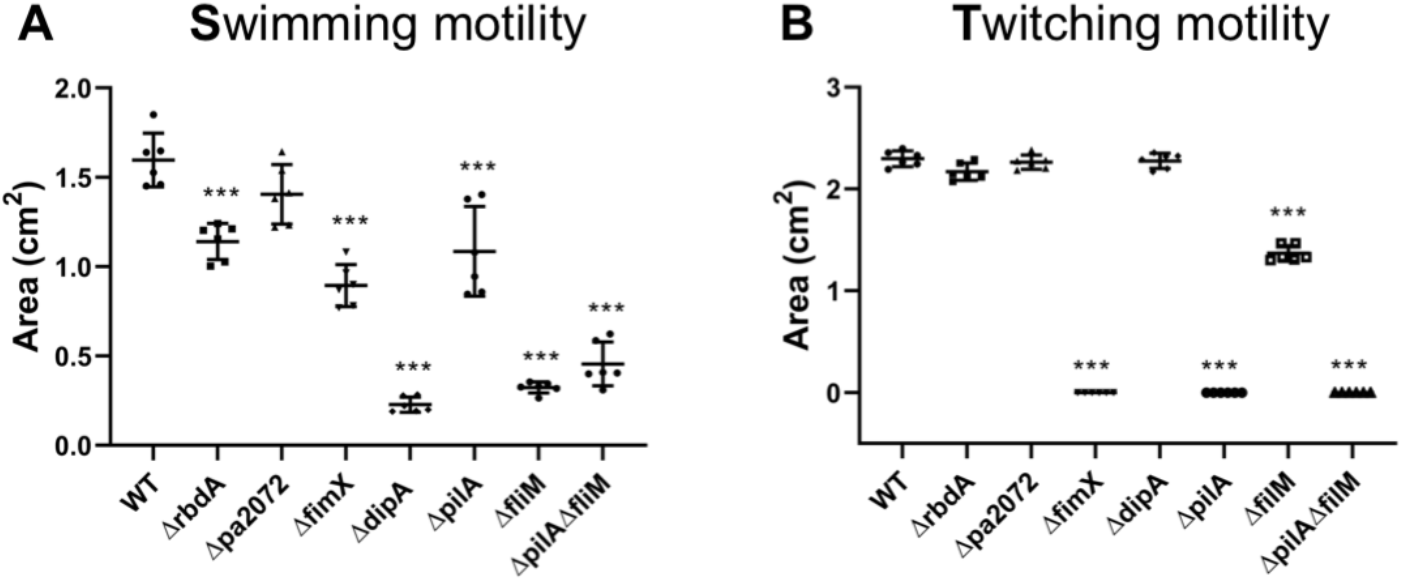
The swimming and twitching phenotypes for deletion mutants. Motilities were determined for the deletion mutants Δ*rbdA*, Δ*pa2072*, Δ*fimX* and Δ*dipA* and compared with the flagellum mutant Δ*fliM*, the pili mutant Δ*pilA*, and the pili flagellum double mutant Δ*pilA*Δ*fliM*. The Student’s T-test was used to determine significances, where *** denotes a confidence level of p<0.01. Acquired from 6 and 10 independent experiments for data shown in (A) and (B), respectively. **(A)** Swimming zones were normalised to PAO1 WT. **(B)** Twitching areas were normalised to PAO1 WT.

impairment that is comparable to the pili mutant Δ*pilA*. In contrast, and as previously reported^14^, pili mediated twitching motility is abolished in *ΔfimX*, similar to the pili *ΔpilA* mutant and the pili flagellum double mutant Δ*pilA*Δ*fliM*, **Figure 4B**.

### Correlation of NO-induced dispersion with c-di-GMP concentration and swarming motility

A NO donor assay was used to explore biofilm dispersal. The efficacies of commercially available donors were determined (see **Table S3** and **Supplement**), and two treatment regimes were selected based on their efficiency to induce dispersal. Generally, a concentration of 250 μM SNP was used for 12 hrs, however for mature biofilms 250 μM S150 was used for 2 hrs to reduce treatment time and side effects observed with SNP (see **Supplement**). Biomass reduction of mature (72 hrs) biofilms grown in MatTek plates was determined, **Figure 5**. Using this metric, NO induced dispersal was impaired in Δ*rbdA*, Δ*pa2072* and Δ*dipA* compared with PAO1 WT, but significantly increased for Δ*fimX*.

**Figure 5.**
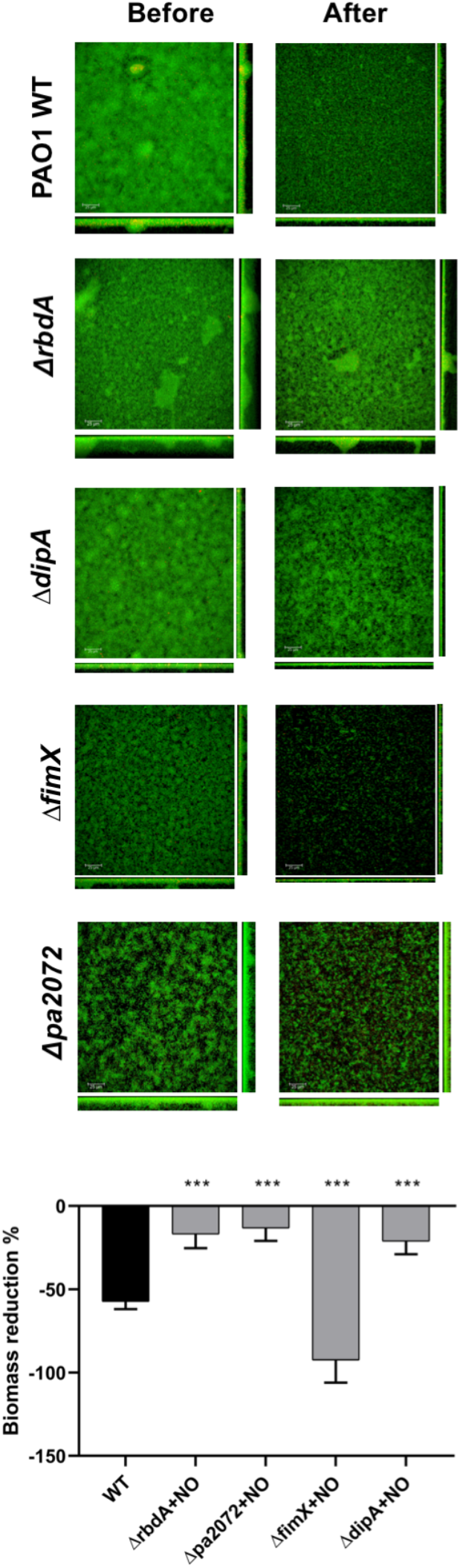
NO-induced dispersal from 72 hrs mature biofilms. Confocal laser scanning microscopy micrographs used for phenotypic analysis of 72 hrs mature biofilms before or after 2 hrs treatment with 250 μM SNP for the deletion mutants, Δ*rbdA*, Δ*pa2072*, Δ*fimX* and Δ*dipA*, compared with PAO1 WT (scale bar 25 μm). The percentage of biofilm mass reduction after treatment was normalised against WT highlighting differences in biomass reduction for the mutants. The Welch T-test was used to determine significances, where *** denotes a confidence level of p<0.01. Data acquired from 3 independent experiments.

As biofilm formation and dispersal are closely linked to intracellular c-di-GMP levels^3^, a c-di-GMP responsive GFP reporter was used to compare relative intracellular levels between each mutant and WT before and after NO treatment. Intracellular c-di-GMP levels were increased within Δ*rbdA* but significantly lowerd for Δ*pa2072*, **Figure 6A**, correlating with their 48 hrs biofilm phenotypes, **Figure 2**. This relationship breaks down for the significantly increased reported c-di-GMP levels of Δ*dipA*, **Figure 6A**, which showed biofilm phenotypes similar to WT, **Figure 2**. Interestingly, all four mutants showed an impaired c-di-GMP response on NO donor addition, implicating them in NO-mediated modulation of intracellular c-di-GMP concentration, **Figure 6B**, despite the impairment of Δ*rbdA*, Δ*pa2072* and Δ*dipA* to disperse, **Figure 5**.

**Figure 6.**
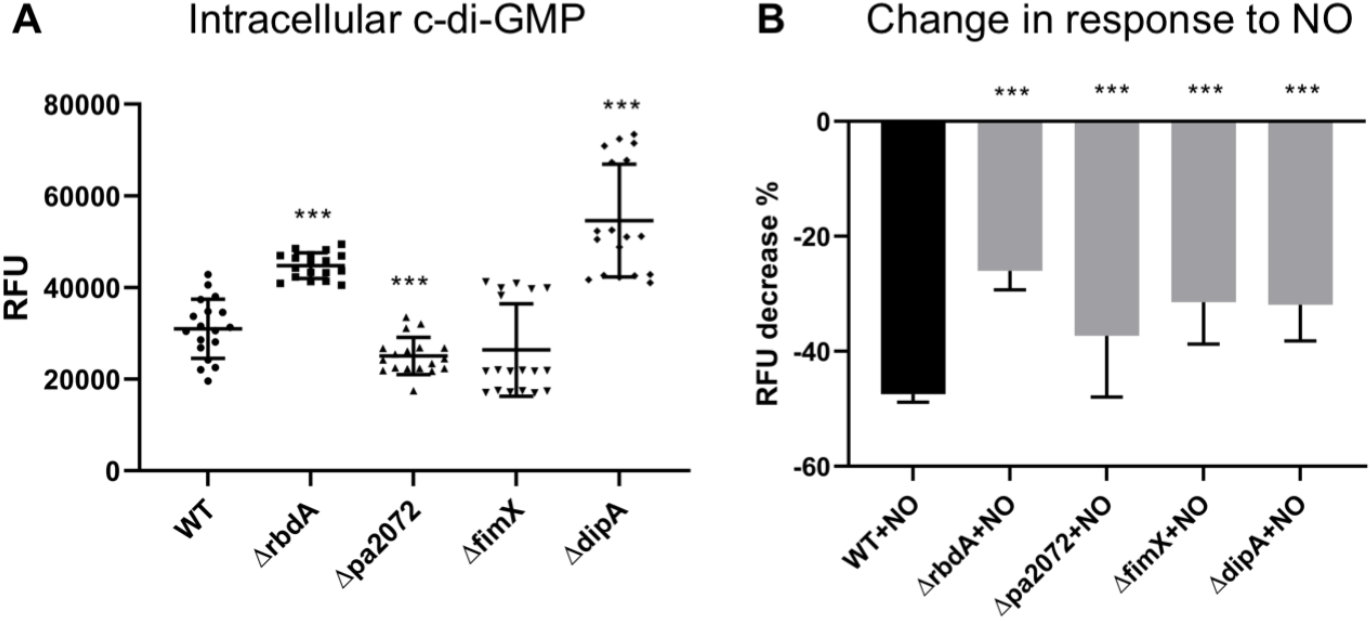
c-di-GMP levels in planktonic culture. The levels of c-di-GMP were quantified using a GFP reporter. **(A)** Relative fluorescence units normalised to PAO1 WT. **(B)** The NO-induced decrease in relative fluorescence, given in percentages. The Student’s T-test was used to determine significances, where *** denotes a confidence level of p<0.01. Data acquired from 3 independent experiments.

We thought to relate the impaired NO dispersal phenotype of Δ*rbdA*, Δ*pa2072* and Δ*dipA* to motility. Biofilm dispersal has been linked to swarming motility, which in *P. aeruginosa* depends on pili and flagella^17,44^, and this motility increases upon exposure to the biofilm dispersal signal NO^45^. Indeed, swarming is strongly supressed in Δ*dipA*, and is also reduced in the presence of NO donor for this mutant, **Figure 7**. Although not as dominant as Δ*dipA*, the deletion mutant Δ*rbdA* also showed a significant reduction in swarming, and swarming is also reduced in the presence of NO donor. The behaviour of Δ*dipA* and Δ*rbdA* is comparable to the flagellum mutant Δ*fliM* and the pili flagellum double mutant Δ*pilA*Δ*fliM*, **Figure 7**. In contrast, Δ*fimX* which dispersed better with the NO donor, **Figure 5**, also showed a larger swarming area than PAO1 WT, comparable to the pili Δ*pilA* mutant, **Figure 7**. Equally, the response to NO donor is significantly more prominent than PAO1 WT, **Figure 7**. Interestingly, Δ*pa2072* showed a swarming phenotype comparable to PAO1 WT and does not display increased swarming upon NO treatment.

**Figure 7.**
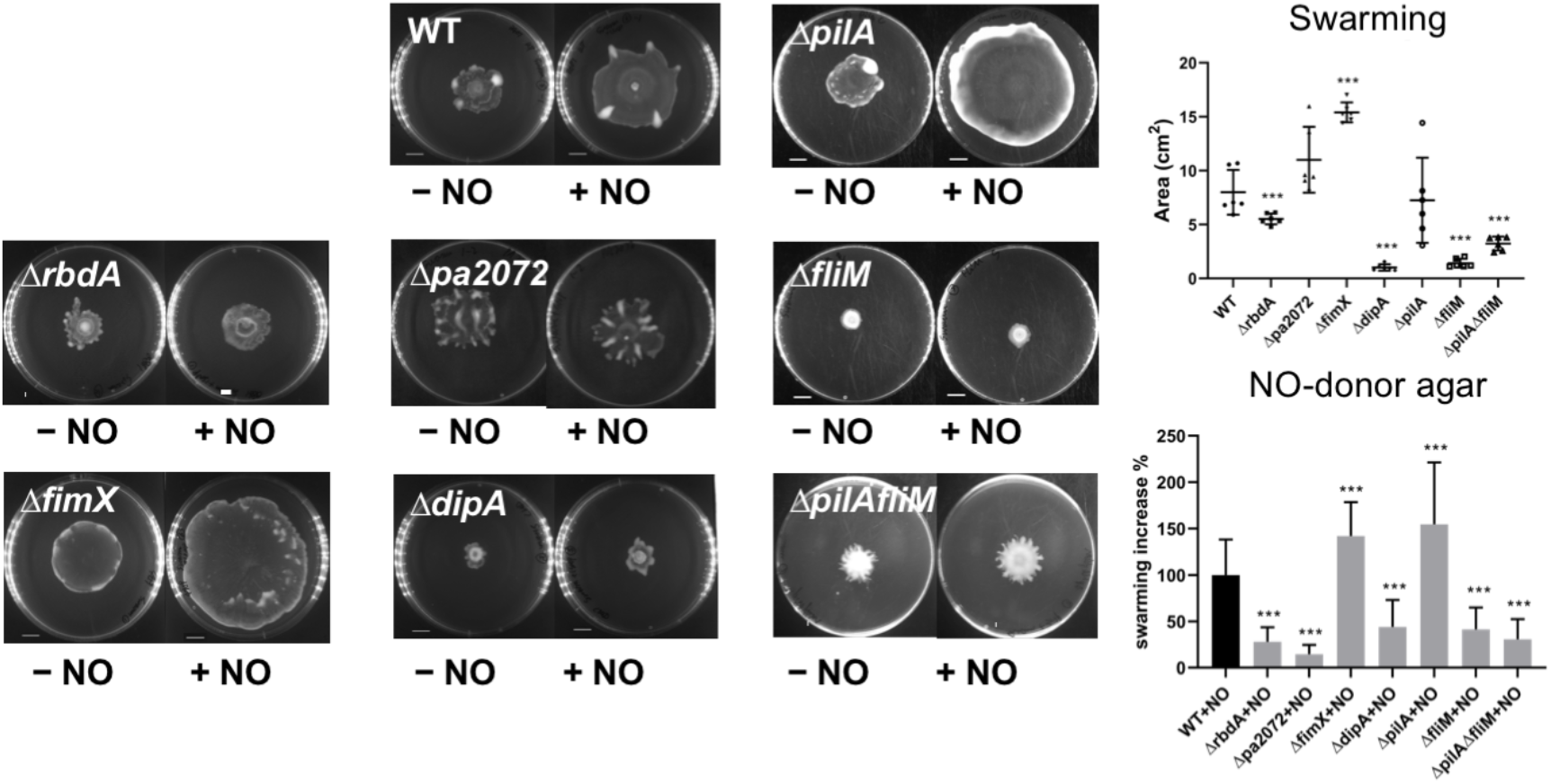
Swarming phenotypes and response to NO dispersal trigger. Motilities were determined for the deletion mutants Δ*rbdA*, Δ*pa2072*, Δ*fimX* and Δ*dipA* and compared with the flagellum mutant Δ*fliM*, the pili mutant Δ*pilA*, and the pili flagellum double mutant Δ*pilA*Δ*fliM*. For plates without NO treatment, swarming areas were normalised to PAO1 WT. The change in swarming areas on NO donor agar (1 μM SNP) was determined and normalised to PAO1 WT. The Student’s T-test was used to determine significances, where *** denotes a confidence level of p<0.01. Data acquired from 6 independent experiments.

Taken together, NO-induced biofilm dispersal and changes in swarming behaviour are correlated for the mutants. For Δ*dipA*, Δ*rbdA* and Δ*pa2072* as well as Δ*fliM* and Δ*fliM*Δ*pilA* swarming is less pronounced or inhibited, **Figure 7**, while NO induced loss in biomass is impaired, **Figure 5**. In contrast, *ΔfimX* as well as Δ*pilA* swarm better than wild type, **Figure 7**, while NO induced loss in biomass is also increased, **Figure 5**.

## Discussion

In *Pseudomonas aeruginosa*, bacterial biofilm formation and dispersal are regulated by intracellular c-di-GMP concentration in response to environmental signals^4,6^. The understanding of enzymes that can regulate c-di-GMP levels, such as diguanylate cyclases (DGCs) and phosphodiesterases (PDEs), and their linked sensor domains will be an essential step to disentangle the network that controls this physiological switch. Bacteria carry these enzymatic activities in multi-domain proteins, allowing for a tight control of catalytic activity and allowing functional diversity through the presence of various regulatory domains. It is unknown whether the total of 38 DGC and PDE enzymes found in *Pseudomonas aeruginosa* PAO1 have redundant functions^5^.

Of the cohort of 38 DGC/PDE proteins, twelve have regulatory PAS domains, **Figure 1**. PAS domains are known as dimerisation domains that respond to a variety of environmental triggers^46,47^. Since DGC and PDE enzymes must dimerise to become catalytically active^3,11,28,48,49^, PAS domains could link a sensory and a dimerisation function and thus provide an elegant solution to integrate signalling with enzyme activation. In the cohort of DGC/PDE proteins, two pairs of highly homologous proteins were identified to test for their functional redundancy. The small proteins PA0290 and PA0338 form the first pair, while PA0861 (RbdA) and PA2072 form another pair. We used a phenotypical selection based on biofilm formation and NO-induced biofilm dispersal, and present data on PA0861 (RbdA), which has been studied before^10,50^, and the previously uncharacterised protein PA2072. Both proteins share a similar structure, with two predicted membrane segments and a cytosolic PAS domain coupled to a DGC/PDE tandem^50^.

Using deletion mutants, we show that both Δ*rbdA* and Δ*pa2072* show differences to WT PAO1 in microcolony formation. The five-fold increase of microcolony size in Δ*rbdA* leads to thicker biofilms and is reflected in a two-fold increase in biomass with enhanced total protein and polysaccharide in the EPS. In stark contrast, microcolony sizes are much reduced for Δ*pa2072*, with consequent formation of much thinner biofilms than WT PAO1 that reached only one fifth of the biomass after 48 hrs. The phenotypes of Δ*rbdA* and Δ*pa2072* also affected biofilm surface coverage, **Figure 2**. The observed phenotypes are in keeping with reported c-di-GMP levels, which are higher for Δ*rbdA* but lower for Δ*pa2072* when compared with PAO1 WT, **Figure 6**. This supports the conclusion that these homologous proteins perform opposite roles, rather than being redundant.

It is notable that several of the 38 *Pseudomonas aeruginosa* DGC and PDE proteins are not enzymes as they show mutations in their catalytic motifs. From the set of twelve PAS containing DGC/PDE proteins, this is true for the well-studied proteins PA4959 (FimX) and PA5017 (DipA). Our *in silico* analysis and earlier data^28,29,31^ indicate FimX has an inactive DGC and supports evidence of an inactive PDE, **Figure 1**. Similarly, while DipA possesses an inactive DGC, its PDE was reported to be active^15^, which is in keeping with the *in silico* analysis performed here. While analysis of biofilm phenotypes showed little difference to PAO1 WT, motilities were severely affected in the deletion mutants, **Figure 4**. Flagellum mediated swimming motility was significantly reduced in Δ*dipA*, as reported previously^18,51^, while *ΔfimX* shows a twitching defect pointing to altered pili function^52^. It is known that FimX is required for the assembly of pili at low intracellular c-di-GMP levels^53^; a demand that can be bypassed when c-di-GMP levels increase^54^. While *ΔfimX* shows comparable c-di-GMP levels to PAO1 WT, deletion of *<u>d</u>ipA* led to an upregulated intracellular c-di-GMP, pointing at a role in c-di-GMP turnover as the protein is an active phosphodiesterase (only the DGC function is perturbed), **Figure 6**. The observations lead to the suggestion that these proteins function in the regulation of motility traits, potentially interacting with other c-di-GMP regulating enzymes through the degenerate DGC (DipA, FimX) or PDE (FimX) domains. This dimerisation behaviour may well be in response to a trigger received by the N-terminal regulatory PAS domains.

The three mutants Δ*rbdA*, Δ*pa2072* and Δ*dipA* highlight a correlation between swarming motility and NO-triggered biofilm dispersal, **Figure 5** and **Figure 7**. Previous work reported a *nirS* mutant unable to produce NO endogenously being deficient in swarming motility, while a nearly 10-fold upregulation of *nirS* was observed in swarming cells compared to non-swarmer planktonic cells^55^. Furthermore, addition of NO was shown not only to decrease intracellular c-di-GMP but also increase swarming in PAO1^45^, suggesting a link between NO, c-di-GMP and swarming regulatory circuits.

We demonstrate decreased swarming for the deletion mutants Δ*rbdA*, Δ*pa2072* and Δ*dipA* and decreased biofilm dispersal, similar to the flagellum and pili/flagellum double deletion mutants Δ*fliM* and Δ*pilA*Δ*fliM*. We note that in Δ*pilA*, the swarming pattern was altered (smooth edges without tendrils). Swarming patterns are similar to the Δ*pilA* mutant for *ΔfimX*, which is in keeping with their abolished pili mediated twitching motility, **Figure 7**. However, increased swarming and better dispersal are documented for Δ*fimX*, similar to the pili mutant Δ*pilA* (for analysis of pili and flagellum deletion biofilm phenotypes see **Figure S1**), further supporting correlation between swarming and NO induced dispersal.

## Conclusion

In this study we demonstrate that PAS containing phosphodiesterases / diguanylate cyclases modulate different facets of motility behaviours, biofilm phenotype and dispersal responses. This is displayed clearly through Δ*rbdA* and Δ*pa2072* gene deletion mutants, which, despite coding for proteins of very similar domain architectures, display almost opposing physiological functions. Studying Δ*fimX* and Δ*dipA*, we add information on pseudo-enzymes into the picture, which prove to play significant regulatory roles not linked to c-di-GMP turnover function. In doing so we highlight an interplay between complex behaviours such as motility and biofilm regulation.

## Acknowledgements

The authors would like to thank Alain Filloux for the generous gift of the modified pCdrA::*gfp* plasmid with tetracycline resistance, Curtis Phippen for help with construction of deletion mutants, Robert Howlin for help with biofilm culture optimisation, David Johnston & Mark Willett for the help with confocal microscopy, Neville Wright for critical discussion of EPS determination. We acknowledge funding by the Diamond Light Source and the University of Southampton to AH.

## Supplement

### NO-donor selection

To date, studies of *P. aeruginosa* biofilm dispersal have utilised a number of different compounds as NO donors, including organic nitrates, nitrite esters, S-nitrosothiols^56^, diazeniumdiolates (NONOates)^57^, and, sodium nitroprusside (SNP)^6,41,45^. Due to this broad range of NO donors, a wide range of concentrations have been reported to be necessary to induce biofilm dispersal from 500 nM to 500 μM^6,15,45,58^. Further complications arise from the chemistry of SNP, which does not release NO without photolysis^59^, or the addition of reducing agents^60^, and in doing so releases the toxic side product cyanide^61^. To determine the most relevant NO donor for this study, we selected a number of commercially available NO donors (Table S3) to quantify their NO release and to test their efficacies to induce PAO1 biofilm dispersal.

#### *NO donor efficacy screening* (modified from O’Toole et al^62^)

An overnight culture of *P. aeruginosa* was diluted into fresh M9 media with an initial inoculum of OD_600nm_ 0.01 and 100μl per well in microtiter plates. The plates were incubated statically at 37°C. For NO treatments, donors listed with their half-life times in Table S3 were tested at concentrations between 1-500 μM, using treatment times between 1-24 hrs. Crystal violet staining showed that 250 μM SNP and 250 μM S150 were sufficient to induce a significant biofilm dispersal (~60%) after 24 and 2 hours, respectively, and these donors were selected for further study **Figure S2A**.

#### *NO release quantification using chemiluminescence* (modified from Piknova et al^40^)

Free NO gas was detected in a chemiluminescence analyser, using an inert carrier gas (synthetic air, 99.99999 % BOC). NO reacts with O_3_ (derived from O_2_ in normal atmospheric conditions) to form nitrogen dioxide (NO_2_) in its activated state, and NO_2_* emits a photon:

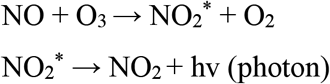

Using a CLD 88Y NO analyser (EcoPhysics, Duernten, Switzerland), the photomultiplier tube was equipped with a long pass filter, detecting emissions above 600 nm, and the NO concentration is calculated from the emitted intensity against a calibration standard^40^. A NO standard was prepared by converting sodium nitrite solution to NO using tri-iodide (I_3_^−^) solution:

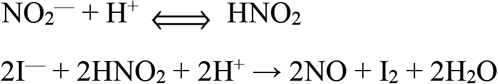

A mixture of KI, I_2_ and glacial acetic acid were used to generate the I_3_ solution. For the I_3_ solution, 500 mg Potassium iodide (KI) and 379.8 mg crystalline iodide (I_2_) were dissolved in 10 ml water to produce final concentrations of 301.2 mM and 137.8 mM respectively. The solution was kept away from light and mixed with glacial acetic acid in a 2:7 ratio by stirring on a magnetic plate for 30 mins. The CLD 88Y was equilibrated at 60 °C starting the carrier gas flow. The I_3_ solution (7.5 ml) was placed into the reaction vessel, bubbling gently through the cell, and NaNO_2_ standards were applied to generate a standard curve (3 replicates each at 250, 375, 500, 750 and 1000 pmol NO). NO donors used a test volume of 9.5 ml (in M9 medium) with 500 μl of stock NO donor solutions at 37 °C. A consistent cold light source was applied as appropriate (Photonic PL^®^3000, maximum light intensity 26 Mlx, color temperature 3250 K). Tracings were recorded at 4 Hz frequency using PowerChrom^®^ (eDAQ Pty LtD, Australia). Data were plotted using Origin 9, calculating areas to quantify NO release, comparing NO donor samples against standards.

We quantified the precise NO release of both SNP and S150 in biofilm culture medium at a concentration of 5 μM across a 2 hrs timeframe, **Figure S2B**. As can be seen from Figure S2B, 50 nmol S150 released NO at a decreasing rate over time and within 1.5 hrs had released around 68.4 nmol NO. SNP showed a constant release of NO in the observation window but much lower than S150, and across 1.5 hours had released 4.69 times less NO than S150. As SNP requires photolysis^60^, we also examined the effect of a cold light source on NO release from SNP. Using 500 μM SNP, we demonstrated that only under strong light induction can SNP release NO efficiently, but this is still much less efficient than the spontaneous donor S150, **Figure S2C**. Therefore, SNP is ideal for long-contact treatment (>12 hrs) under normal light conditions, while S150 is useful for short-contact treatment (2 hrs), according to specific requirements in different experiments. As such, SNP was chosen for 24 hrs incubation, used for swarming agar assays, while S150 was used for triggering mature biofilm dispersal within 2 hrs contact time.

**Table S1:** Bacterial strains and plasmids

| Strains and plasmids | Genotype or phenotypes<sup>a</sup> | Reference |
| --- | --- | --- |
| *E. coli* |  |  |
| S17-1(λpir) | *TpR SmR recA, thi, pro, hsdR-M+RP4; 2-Tc:Mu; Km Tn7* | Biomedal corp. |
| DH5α | *spuE44 ΔlacU169(φ80lacZΔM15) hsdR17 λpir recA1 endA1 gyrA96 thi-1 relA1* | Lab collection |
| *P. aeruginosa* |  |  |
| PAO1 | Wild-Type, C.Manoil lab, University of Washington | 63 |
| **KO mutants** |  |  |
| Δpa0285 | PAO1 mutant (*pa0285::aacCI*); Gm<sup>R</sup> | This study |
| Δpa0290 | PAO1 mutant (*pa0290::aacCI*); Gm<sup>R</sup> | This study |
| Δpa0338 | PAO1 mutant (*pa0338::aacCI*); Gm<sup>R</sup> | This study |
| Δpa0575 | PAO1 mutant (*pa0575::aacCI*); Gm<sup>R</sup> | This study |
| Δpa0847 | PAO1 mutant (*pa0847::aacCI*); Gm<sup>R</sup> | This study |
| Δpa0861 (*ΔrbdA*) | PAO1 mutant (*pa0861::aacCI*); Gm<sup>R</sup> | This study |
| Δpa1181 (*ΔyegE*) | PAO1 mutant (*pa1181::aacCI*); Gm<sup>R</sup> | This study |
| Δpa2072 | PAO1 mutant (*pa2072::aacCI*); Gm<sup>R</sup> | This study |
| Δpa4601 (*ΔmorA*) | PAO1 mutant (*pa4601::aacCI*); Gm<sup>R</sup> | This study |
| Δpa4959 (*ΔfimX*) | PAO1 mutant (*pa4959::aacCI*); Gm<sup>R</sup> | This study |
| Δpa5017 (*ΔdipA*) | PAO1 mutant (*pa5017::aacCI*); Gm<sup>R</sup> | This study |
| Δpa5442 | PAO1 mutant (*pa5442::aacCI*); Gm<sup>R</sup> | This study |
| Δpa1443 (*ΔfliM*) | PAO1 mutant (*pa4959::aacCI*); Gm<sup>R</sup> | This study |
| Δpa4525 (*ΔpilA*) | PAO1 mutant (*pa5017::aacCI*); Gm<sup>R</sup> | This study |
| Δpa1443Δpa4525 (*ΔpilAΔfliM*) | PAO1 mutant (*pa1443::aacCI, pa4525::Neo*); Gm<sup>R</sup>; Km<sup>R</sup> | This study |
| **KO mutants with c-di-GMP reporter gfp vector** |  |  |
| PAO1 Tn7CdrA::gfp | PAO1 WT with mini-Tn7-P<sub>cdrA</sub>-RBSII-gfp(Mut3)-T<sub>0</sub>-T<sub>1</sub>; Tet<sup>R</sup> | This study |
| Δpa0285 Tn7CdrA::gfp | PAO1 mutant (*pa0285::aacCI*) with mini-Tn7-P<sub>cdrA</sub>-RBSII-gfp(Mut3)-T<sub>0</sub>-T<sub>1</sub>; Gm<sup>R</sup>; Tet<sup>R</sup> | This study |
| Δpa0290 Tn7CdrA::gfp | PAO1 mutant (*pa0290::aacCI*) with mini-Tn7-P<sub>cdrA</sub>-RBSII-gfp(Mut3)-T<sub>0</sub>-T<sub>1</sub>; Gm<sup>R</sup>; Tet<sup>R</sup> | This study |
| Δpa0338 Tn7CdrA::gfp | PAO1 mutant (*pa0338::aacCI*) with mini-Tn7-P<sub>cdrA</sub>-RBSII-gfp(Mut3)-T<sub>0</sub>-T<sub>1</sub>; Gm<sup>R</sup>; Tet<sup>R</sup> | This study |
| Δpa0575 Tn7CdrA::gfp | PAO1 mutant (*pa0575::aacCI*) with mini-Tn7-P<sub>cdrA</sub>-RBSII-gfp(Mut3)-T<sub>0</sub>-T<sub>1</sub>; Gm<sup>R</sup>; Tet<sup>R</sup> | This study |
| Δpa0847 Tn7CdrA::gfp | PAO1 mutant (*pa0847::aacCI*) with mini-Tn7-P<sub>cdrA</sub>-RBSII-gfp(Mut3)-T<sub>0</sub>-T<sub>1</sub>; Gm<sup>R</sup>; Tet<sup>R</sup> | This study |
| Δpa0861 Tn7CdrA::gfp | PAO1 mutant (*pa0861::aacCI*) with mini-Tn7-P<sub>cdrA</sub>-RBSII-gfp(Mut3)-T<sub>0</sub>-T<sub>1</sub>; Gm<sup>R</sup>; Tet<sup>R</sup> | This study |
| Δpa1181 Tn7CdrA::gfp | PAO1 mutant (*pa1181::aacCI*) with mini-Tn7-P<sub>cdrA</sub>-RBSII-gfp(Mut3)-T<sub>0</sub>-T<sub>1</sub>; Gm<sup>R</sup>; Tet<sup>R</sup> | This study |
| Δpa2072 Tn7CdrA::gfp | PAO1 mutant (*pa2072::aacCI*) with mini-Tn7-P<sub>cdrA</sub>-RBSII-gfp(Mut3)-T<sub>0</sub>-T<sub>1</sub>; Gm<sup>R</sup>; Tet<sup>R</sup> | This study |
| Δpa4601 Tn7CdrA::gfp | PAO1 mutant (*pa4601::aacCI*) with mini-Tn7-P<sub>cdrA</sub>-RBSII-gfp(Mut3)-T<sub>0</sub>-T<sub>1</sub>; Gm<sup>R</sup>; Tet<sup>R</sup> | This study |
| Δpa4959 Tn7CdrA::gfp | PAO1 mutant (*pa4959::aacCI*) with mini-Tn7-P<sub>cdrA</sub>-RBSII-gfp(Mut3)-T<sub>0</sub>-T<sub>1</sub>; Gm<sup>R</sup>; Tet<sup>R</sup> | This study |
| Δpa5017 Tn7CdrA::gfp | PAO1 mutant (*pa5017::aacCI*) with mini-Tn7-P<sub>cdrA</sub>-RBSII-gfp(Mut3)-T<sub>0</sub>-T<sub>1</sub>; Gm<sup>R</sup>; Tet<sup>R</sup> | This study |
| Δpa5442 Tn7CdrA::gfp | PAO1 mutant (*pa5442::aacCI*) with mini-Tn7-P<sub>cdrA</sub>-RBSII-gfp(Mut3)-T<sub>0</sub>-T<sub>1</sub>; Gm<sup>R</sup>; Tet<sup>R</sup> | This study |
| **Plasmids** |  |  |
| pPS856 | Ap<sup>R</sup>, Gm<sup>R</sup>; 0.83-kb blunt-ended SacI fragment from pUCGM | 37 |
| pEX100T | Ap<sup>R</sup>; oriT<sup>+</sup> sacB<sup>+</sup> gene replacement vector | 37 |
| pCdrA::gfp | pUCP22Not-P<sub>cdrA</sub>-RBSII-gfp(Mut3)-T<sub>0</sub>-T<sub>1</sub>, Amp<sup>R</sup> Tet<sup>R</sup> | 42 |
| pCR<sup>TM</sup>4-TOPO<sup>TM</sup> | Km<sup>R</sup> and Amp<sup>R</sup> TOPO cloning expression vector | Invitrogen<sup>TM</sup> |

**Table S2:**
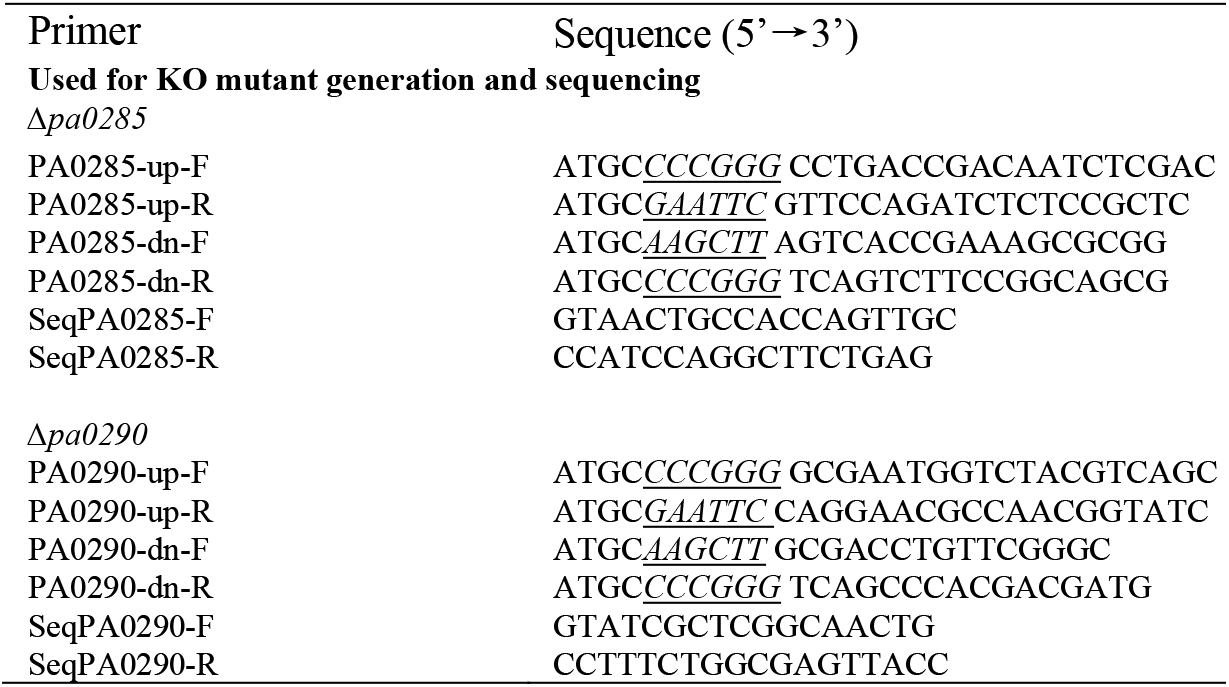
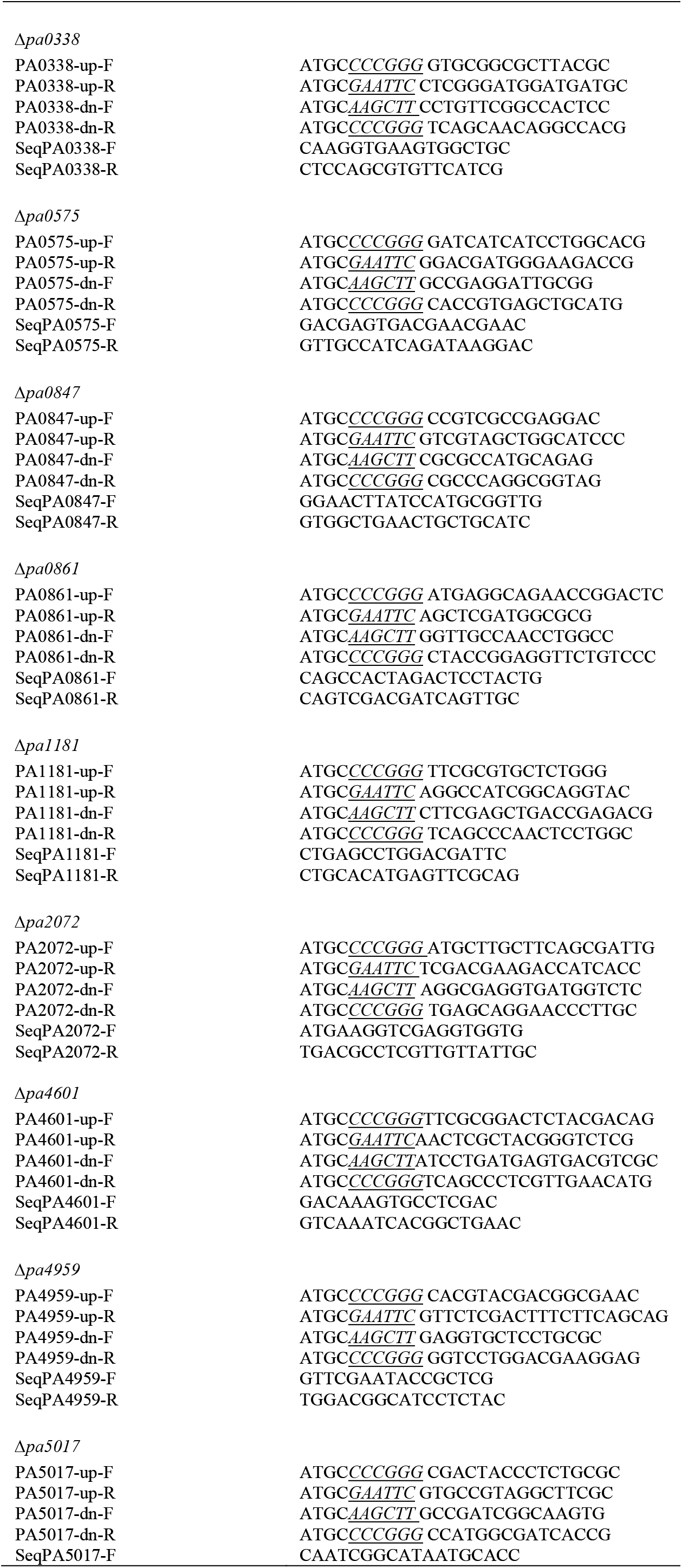
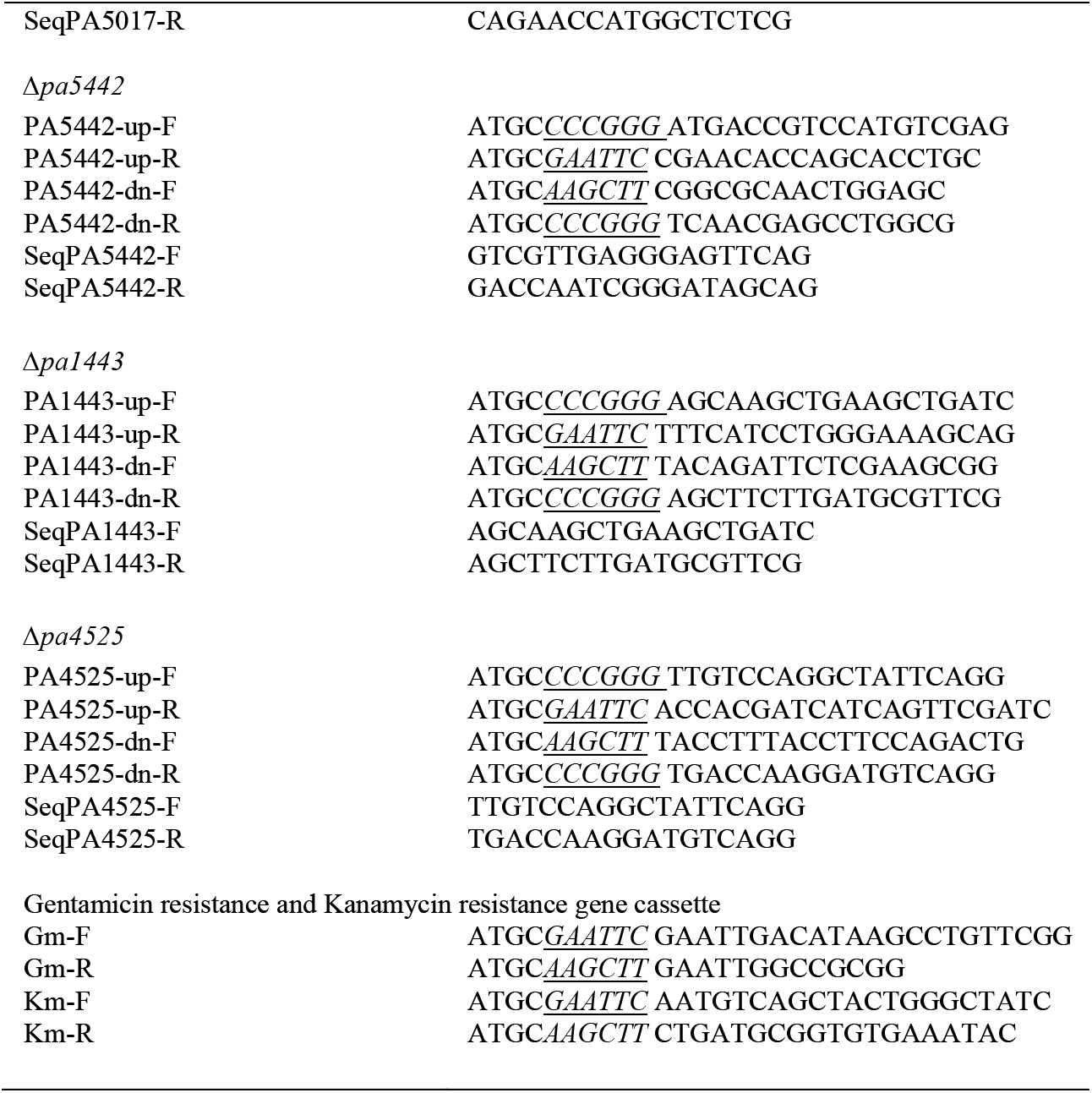
Primer sequences

**Table S3:** commercially available NO donors. Half-live of seven commercially available NO donors at 37 °C, pH 7.4.

|  |  |
| --- | --- |
| SNP | <2 mins |
| SNAP | 6 hrs |
| GSNO | 1-3 hrs |
| PROLI NONOate | 1.8 secs |
| MAHMA NONOate (NOC-9) | 1 min |
| Spermine NONOate (S150) | 39 mins |
| DEA-NONOate | 2 mins |

**Figure S1.**
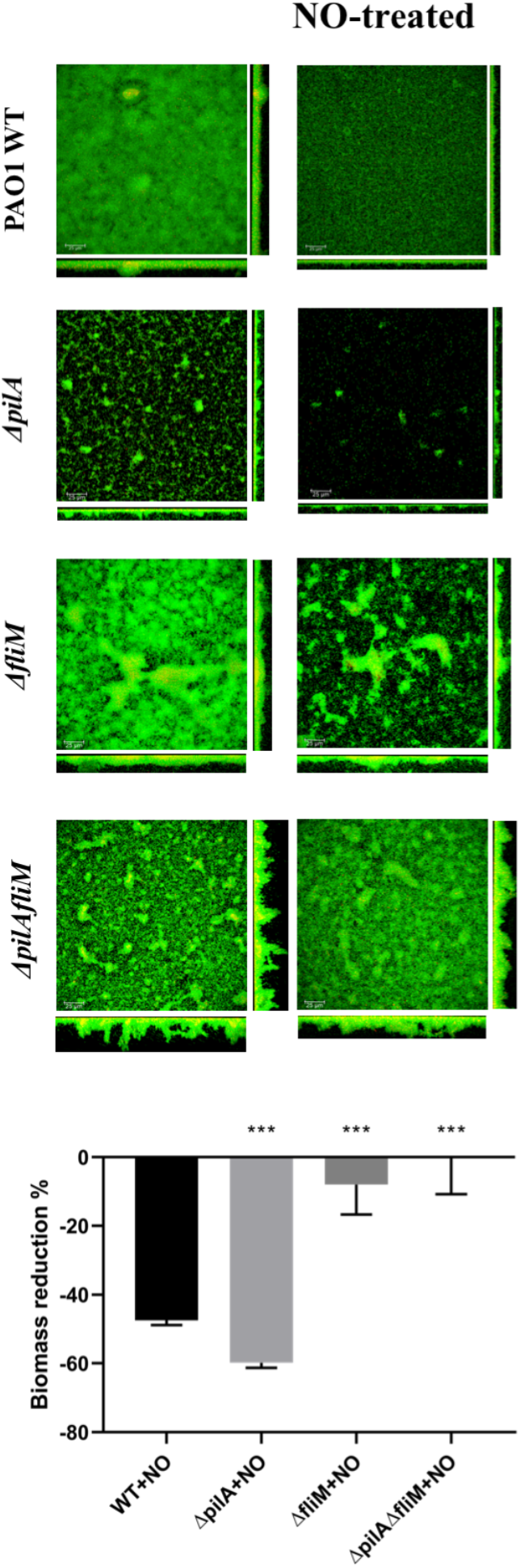
NO-induced biofilm dispersal of flagellum and pili deletion mutants. Confocal laser scanning microscopy images used for phenotypic analysis of 72 hrs mature biofilms before or after 2 hrs treatment with 250 mM SNP for the flagellum mutant Δ*fliM*, the pili mutant Δ*pilA* and the double mutant Δ*pilA*Δ*fliM* compared with PAO1 WT (scale bar 25 μm). Biofilm mass after treatment was normalised against WT, highlighting differences in biomass reduction for the mutants. The Student’s T-test was used to determine significances, where *** denotes a confidence level of p<0.01, and ** denotes a confidence level of 0.01<p<0.05. Data acquired from 3 independent experiments.

**Figure S2.**
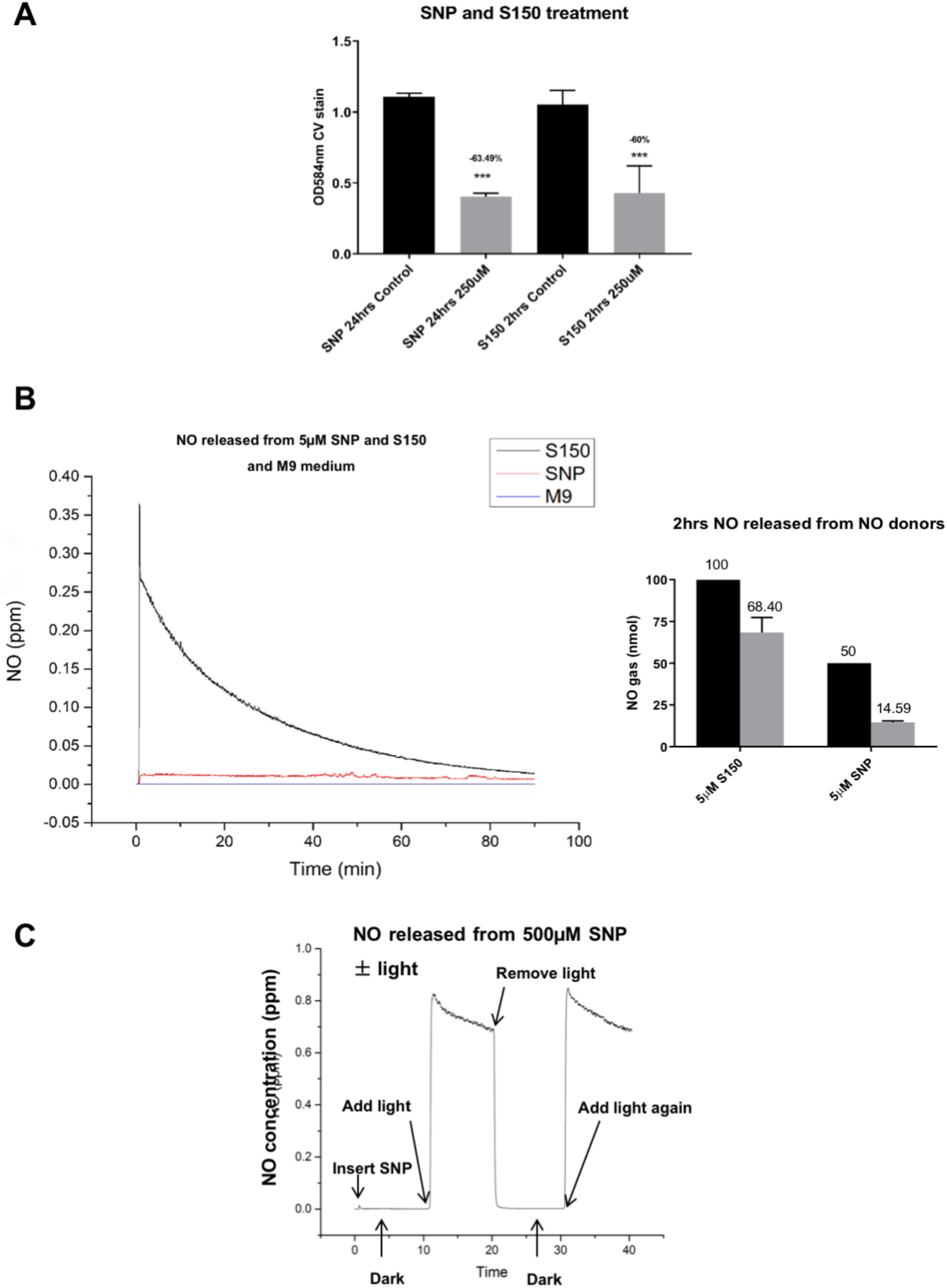
Comparison of S150 and SNP NO donors. **(A)** The efficiencies of SNP and S150 on triggering PAO1 WT biofilm dispersal. Data represent data means of n=6 of 3 biological replicates, with ** denoting 0.01<p<0.05 and *** denoting p<0.01. **(B)** NO release curves using 5 μM S150, 5 μM SNP or M9 media (control). Inset: the amount of NO released. The Student’s T-test was used to determine significances, where ** denotes a confidence level of 0.01<p<0.05, and *** denotes a confidence level of p<0.01. Data acquired from 3 independent experiments. **(C)** NO release curve from 500 μM SNP with/without cold light source at 37 °C in M9 media.

